# Prober: A general toolkit for analyzing sequencing-based ‘toeprinting’ assays

**DOI:** 10.1101/063107

**Authors:** Bo Li, Akshay Tambe, Sharon Aviran, Lior Pachter

## Abstract

A number of high-throughput transcriptase drop-off assays have recently been developed to probe post-transcriptional dynamics of RNA-protein interaction, RNA structure, and post-transcriptional modifications. Although these assays survey a diverse set of ‘epitranscriptomic’ marks, they share methodological similarities and as such their interpretation is predicated on addressing similar computational challenges. Among these, a key question is how to learn isoform-specific chemical modification profiles in the face of complex read multi-mapping. In this paper, we propose PROBer, the first rigorous statistical model to handle these challenges for a general set of sequencing-based ‘toeprinting’ assays.

## INTRODUCTION

While much of the control of gene expression occurs via transcriptional regulation, it is becoming increasingly clear that post-transcriptional regulation also plays a key role in modulating expression products (Schwanhäusser et al., 2011). Several mechanisms contribute to this phenomenon, including covalent posttranscriptional chemical modification of RNA molecules (Roundtree and He, 2016), protein binding and the assembly of higher-order ribonucleoprotein complexes (Glisovic et al., 2008), and the ability of RNA molecules to fold into and switch between intricate 2- and 3- dimensional folds (Mortimer et al., 2014; Schwanhäusser et al., 2011; Wan et al., 2011). Understanding both the expression level and the ‘meta-information’ (post-transcriptional marks) associated with a given transcript can shed light not only on the functions that an individual sequence performs, but also on the cellular pathways that it participates in and controls.

Recent advances in massively parallel DNA sequencing have enabled the transcriptome-wide investigation of several ‘epitranscriptomic’ layers. Although the specific of the assays differ depending on the specific chemicals used, there are several that share a common theme. We term these experiments ‘toeprinting’ (Hartz et al., 1988) by high-throughput sequencing (Figure 1A) as they share a common workflow: chemically modifying RNAs to encode a signal of interest, decoding these chemical signals by reverse transcriptase drop-off, and lastly, sequencing and mapping the resulting cDNA toeprints to recover the chemical modification ‘signatures’.

**Figure 1.**
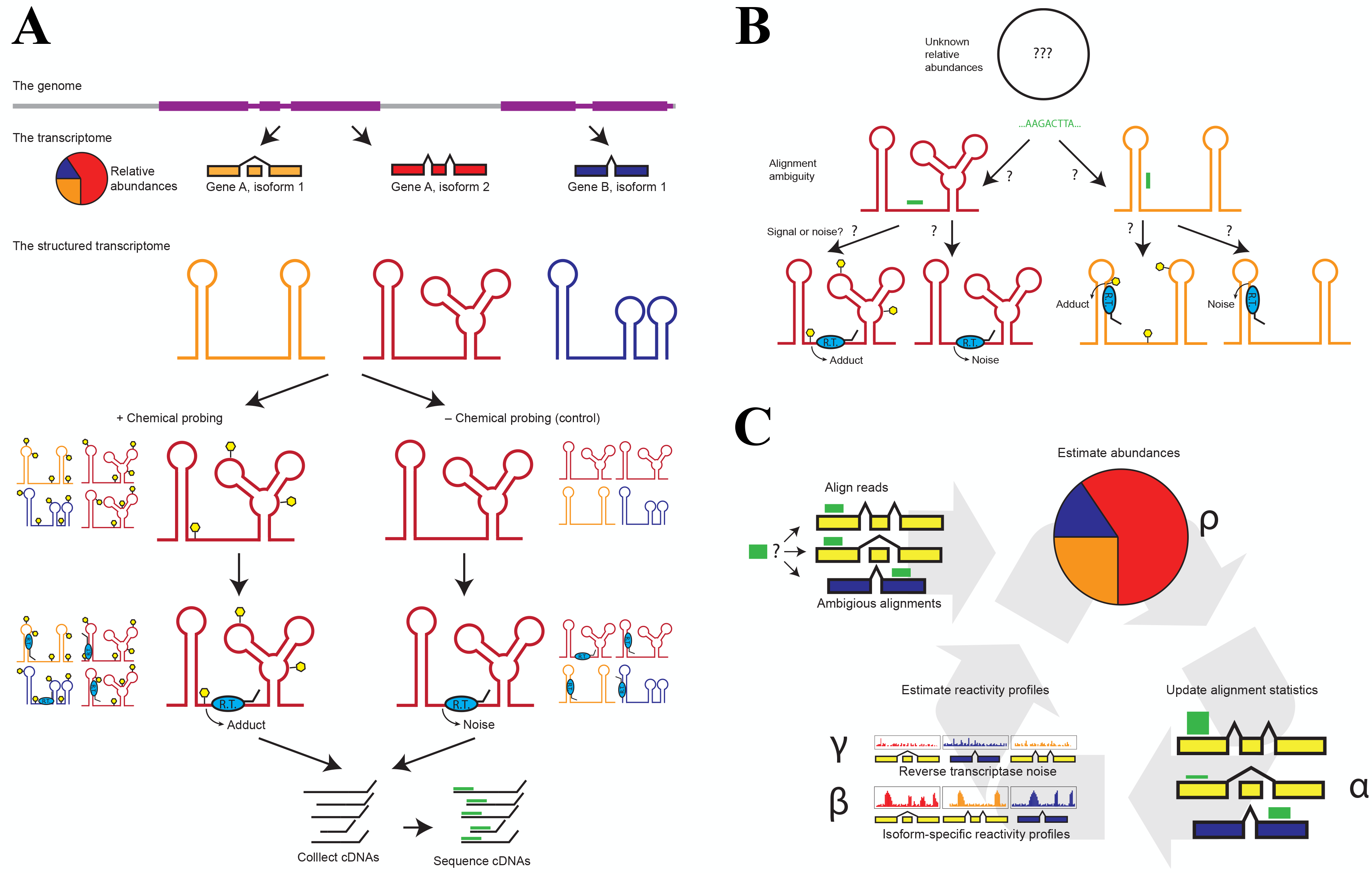
Cartoon depictions of sequencing-based “toeprinting” experiments, the associated Bioinformatics challenges, and our solution. (A) Cartoon depiction of an idealized toeprinting experiment. The genome is transcribed and RNAs are spliced and folded to form the structured transcriptome. This pool of RNAs is split into two, and either treated with a chemical probe, or mock-treated without the chemical probe. These chemical adducts are detected by reverse transcriptase (RT) drop-off, but the signal is convoluted by reverse transcriptase noise. Reverse transcription products are collected and sequenced. (B) Potential bioinformatics challenges. The structured transcriptome that gave rise to a given toeprinting dataset consists of known transcripts of unknown relative abundance. Reads from this dataset might align ambiguously to one or more transcripts, and might have been generated by either RT drop-off at a chemical modification, or by RT noise. (C) Conceptual workflow of PROBer. Sequencing data (both treatment and control datasets) from a toeprinting experiment are used as the input. In the E- step, reads are assigned to transcripts depending on an initial alignment, and the relative abundances & toeprinting parameters of the transcripts estimated in the M-step. In the M- step, transcript abundances and toeprinting parameters are learned, using the read assignments calculated in the E-step.

Within this framework, the iCLIP protocol (König et al., 2010) explores RNA-protein interactions, SHAPE and DMS probing (Ding et al., 2014; Rouskin et al., 2014; Spitale et al., 2015; Talkish et al., 2014) explore RNA secondary structure by using selective chemical probes to modify and ‘emark’ unpaired flexible nucleotides, and Pseudo-seq (Carlile et al., 2014) detects RNA pseudouridylation by utilizing a reagent which specifically forms adducts at pseudouridine sites (Ψs). In each of these experiments, the upstream chemical modification is widely variable, but the library preparation and sequencing techniques are essentially the same: reverse transcription in a manner where cDNAs preferentially terminate at the sites of chemical modification, adaptor ligation to the site of reverse transcriptase drop-off, and PCR amplification followed by sequencing of the cDNA resulting library. Additionally, the number of characterizable epitranscriptomic marks is ever expanding, as are the associated chemical toolkits. As a result, ‘toeprinting’, by high-throughput sequencing is becoming an essential tool for probing post-transcriptional regulation.

A key step in analyzing ‘toeprinting’ experiments is to accurately learn reverse transcriptase drop-off profiles from the sequence data. These profiles are subsequently used to infer, for example, sequence motifs, secondary structure predictions, or sites of posttranscriptional chemical modification. Data produced in the experiments potentially contain multiple layers of valuable information: reads contain information about both modification at sites as well as about the identity and abundance of RNA transcripts. The ability to make full use of this information becomes the key for accurate estimation of drop-off profiles and requires addressing associated bioinformatics challenges including the conflation of read counts by reverse transcriptase noise, variable transcript abundances, and read mapping ambiguity. However, to date, the proposed solutions to these problems have mainly consisted of ad-hoc heuristics rather than statistical modeling.

## RESULTS

### Bioinformatics challenges

Accurately determining the transcript abundances and drop-off profiles in ‘toeprinting’ experiments is complicated by several factors (Figure 1B) (Aviran and Pachter, 2014). Such experiments face a problem that is fundamental in RNA-seq: reads align ambiguously to multiple transcripts, and appropriately handling ambiguously mapped reads (which can represent a significant proportion of alignable reads in such experiments, see Table S1 and S2) is imperative to correctly learning transcript abundances (Bray et al., 2016; Li and Dewey, 2011; Li et al., 2010; Roberts and Pachter, 2013; Trapnell et al., 2010). Incorrectly allocating multi-mapping reads adversely affects the estimated abundances of not only the transcripts that the reads were misallocated to/from, but also abundance estimates of related transcripts.

In toeprinting experiments, the multi-mapping problem is further exacerbated by the fact that accurate estimation of the RNA chemical modification probabilities depends on both correctly allocating multimapped reads, and deconvolving chemical modification profiles from adduct-independent noisy RT drop-off. All of these factors are inter-related and poor estimation of any one of them may significantly skew estimates of the others. Yet all of these factors must be accounted for to quantitative estimate modification rates.

### Our solution: the PROBer software

To address the computational challenges associated to the interpretation and analysis of toeprinting assays we have developed a statistically rigorous approach that serves the dual purpose of unifying these assays via a shared computational framework, while providing an approach to inference that is robust to small variances in experimental protocol. Our methods are implemented in software, termed PROBer, that is based on a statistical model to jointly infer transcript abundance and modification probabilities, as well as several other parameters (see Experimental Procedures and Figure S1A) and was developed by building on previous work on RNA-Seq (Bray et al., 2016; Li et al., 2010; Li and Dewey, 2011; Roberts and Pachter, 2013; Trapnell et al., 2010), as well as models for simpler single-transcript structure-probing SHAPE-seq experiments (Aviran et al., 2011a; 2011b). The PROBer model assumes that the input data consists of raw reads (either single- or paired- end) obtained separately from a chemically treated sample, containing information about modification probabilities, and from a mock-treated control, informing about noise parameters. It assumes that cDNA fragments were generated by first selecting a transcript from the transcriptome (according to its abundance and length), randomly priming (or fragmenting) that transcript, and primer extending one nucleotide at a time. At each nucleotide encountered by the reverse transcriptase (RT) in this process, there is some probability of terminating the reverse transcription, due to modification, RT noise, primer collision, or encountering the end of the template fragment. A cDNA fragment generated by this process is observed as sequenced read if it passes a size-selection filter, which is dependent on the fragment length. From this the extent to which all the parameters in the experiment are inter-related becomes clear.

We implemented an Expectation-Maximization algorithm (Dempster et al., 1977) in PROBer to infer the parameters of the model (see Experimental Procedures). As in many cases it is of interest to have transcript-specific modification profiles rather than genes; indeed for structure probing experiments it is meaningless to consider the modification profiles of a gene, so we modeled modification at the isoform level. The PROBer workflow, shown schematically in Figure 1C, begins with a set of reads alignments (separately for the chemically-treated experiment and the untreated control). Starting with initial parameter estimates, reads are allocated to transcripts based on both abundance and structure parameters. The allocated read ‘pseudocounts’ are then used to estimate *maximum a posterior* (MAP) modification probabilities as well as RT noise, size-selection terms and maximum-likelihood estimates (MLEs) of transcript abundances. These steps are repeated until convergence.

### PROBer outperforms other methods in profiling RNA structures

To test the accuracy of PROBer on structure-probing experiments we investigated its performance on both simulated and experimental data. In simulations, we generated a dataset in a manner consistent with the chemical mapping protocol (see Experimental Procedures) and attempted to recover parameter estimates from these simulated reads alone. As shown in Figure 2A, at a global scale, PROBer yielded significantly improved parameter estimates when compared with other approaches. These results were representative of multiple simulations (Figure S3C) and this improvement was observed across a range of expression levels. In addition, because PROBer takes structure information into consideration, it also estimates transcript abundances better than popular RNA-Seq quantification tools that are not aware of RNA structures (Figure S4).

**Figure 2.**
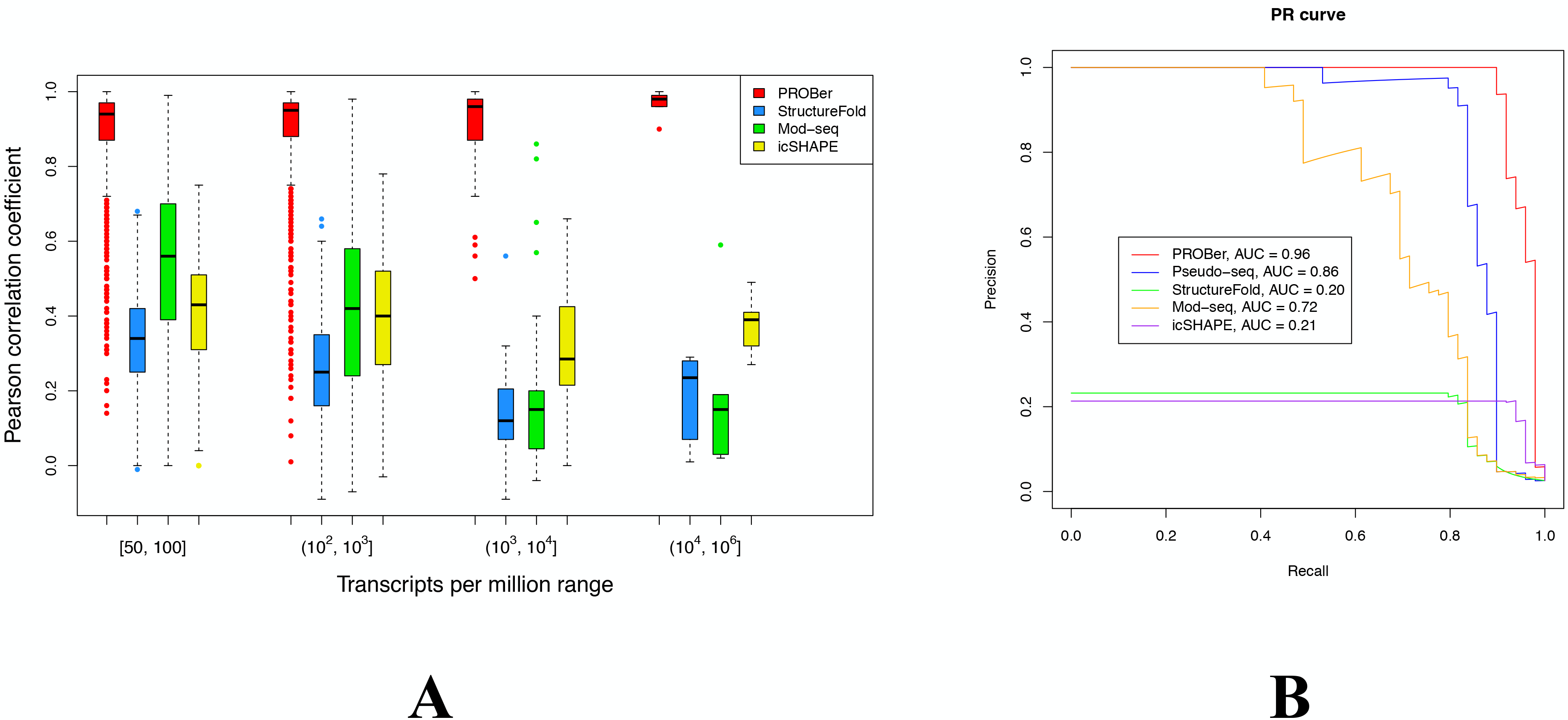
Performance of PROBer as compared to alternative approaches. (A) A simulated RNA structure-probing dataset was generated in a manner consistent with Ding et al. (Ding et al., 2014), and used as the input for a number of structure-probing quantification methods, which include PROBer, StructureFold (Tang et al., 2015), Mod-seq (Talkish et al., 2014), and icSHAPE (Spitale et al., 2015). Accuracy was evaluated by comparing the results from these methods to the simulation parameters using Pearson’s correlation coefficient. PROBer consistently outperforms all other methods across a wide range of expression levels. See also Figure S3C and Experimental Procedures. (B) PROBer was compared to alternative methods on data for predicting known pseudouridine (Ψ) sites in yeast rRNAs and snoRNA (Carlile et al., 2014). Methods were evaluated by Precision-Recall (PR) curves and area under curve (AUC) values. PROBer outperforms all other methods significantly. See also Figure S6 and S7.

PROBer’s performance at recovering secondary structure constraints for transcripts with moderate expression levels (between 100 and 1000 TPM) vastly improves on alternative methods at the highest expression levels (greater than 10,000 TPM). This result indicates that PROBer requires approximately 90% less data (when compared to ad hoc methods) to produce structural estimates of equal or better accuracy. As transcript abundances follow an exponential distribution, a moderate improvement in the range of expression levels that yields useful structural constraints translates to a large increase in the number of transcripts that can be probed. Thus, PROBer allows the experimenter to access a larger fraction of the transcriptome at the same sequencing depth and experimental cost.

As these simulations were based on the same generative model for structure probing experiments that the PROBer software uses, we were concerned that our protocol would artificially inflate the apparent performance of PROBer. We therefore included in our simulated transcriptome three additional transcripts for which chemical modification profiles have been independently measured by SHAPE-seq (Aviran et al., 2011b; Lucks et al., 2011). These transcripts (see Experimental Procedures) served as a “digital spike-in”, allowing us to verify that our simulated experiments were not biased by the simulation parameters. The accuracy of PROBer was confirmed by these digital spike-in experiments (Table S3).

We further tested whether this improvement was also evident in real datasets by examining modification probability estimates for ribosomal RNAs, which have well-characterized structures (Cannone et al., 2002). We calculated receiver operating characteristic (ROC) curves on a variety of structure-probing data sets (Ding et al., 2014; Spitale et al., 2015; Talkish et al., 2014) (Figure S5). These data sets were from different organisms (Arabidopsis, yeast, or mouse), used different chemical agents (DMS or NAI-N3), and adopted different priming methods (random priming or fragmentation). While performing this analysis we observed that all methods tested showed similar area under ROC curve (AUROC) values, indicating that DMS reactivity might not be captured by the consensus secondary structure ground-truth.

### PROBer identifies more true Ψs than other methods

As the lack of other experimentally validated secondary structures prevented us from further studying the performance of PROBer in structure profiling, we therefore explored whether it could be used to quantify pseudouridylation profiles produced by CMCT modification of RNA. As this experimental method performs the RNA modification reaction in purified (and likely denatured / unfolded) RNA, we did not expect solvent accessibility to confound the chemical modification signal. We analyzed Pseudo-seq data from (Carlile et al., 2014) and used all known Ψ sites in ribosomal and small nuclear RNAs as a ground truth, to which we compared PROBer estimated modification profiles. Precision-recall curve analysis of these data revealed that PROBer outperforms existing ad hoc methods for predicting Ψ. Importantly, PROBer was able to detect an experimentally validated pseudo-U site (m^1^acp^3^Ψ1191 in 18S rRNA) that was not detected by ad hoc approaches (Figure S6). This indicates that PROBer is capable of capturing biologically relevant information that would be otherwise lost.

### PROBer detects more putative protein binding sites with canonical motifs

Next, we tested PROBer on iCLIP data. The iCLIP experiment encodes protein binding information in a toeprinting-type manner by crosslinking RNA to proteins and degrading the crosslinked protein by proteolysis. This leaves a short peptide fragment attached to the site on the RNA where it was crosslinked, and that can therefore cause RT drop-off.

The iCLIP protocol differs from other ‘toeprinting’ protocols in two aspects: First, the RNase degradation process produces fragments that are only around the crosslink sites. This results in sparse iCLIP read alignment to the genome. Second, it does not include a sequenced control that can help reduce background noise. Therefore, PROBer uses a simpler model (Figure S1B) to allocate ambiguously mapped iCLIP reads, which compose a significant portion of iCLIP data (Table S2).

We reanalyzed one set of iCLIP data generated by Nostrand et al. (Van Nostrand et al., 2016), which investigated the transcriptome-wide binding preferences for the RNA regulating protein RBFOX2. Importantly, the UGCAUG binding motif of this protein has been validated *in vitro*, providing an independent ground truth for our evaluation.

As expected, our analysis of this dataset with PROBer yielded the sequence motif that was previously reported, indicating that the PROBer model can indeed handle iCLIP data as well. More importantly, we found that PROBer can detect extra binding sites with the exact sequence motif, which could not be detected if we restrict to uniquely mapping reads. Figure 3A gives an example in protein-coding gene *NUP133*. The detected binding site is located at the intronic region downstream of exon 15 of *NUP133*, which implies RBFOX2 may promote the inclusion of this exon in the mature transcripts (Yeo et al., 2009). Note that this significantly enriched binding site cannot be detected using only uniquely mapping reads. We compared PROBer with a baseline method that distributes multi-mapping reads evenly to all aligned locations. Superior performance of PROBer over the baseline method (Figure 3C) demonstrated the power of PROBer’s statistically sound iCLIP model. Our results with the RNA binding protein RBFOX2 clearly demonstrate that multi-mapping iCLIP reads contain valuable information and that the common practice of restricting analysis to unique mappings is suboptimal.

**Figure 3.**
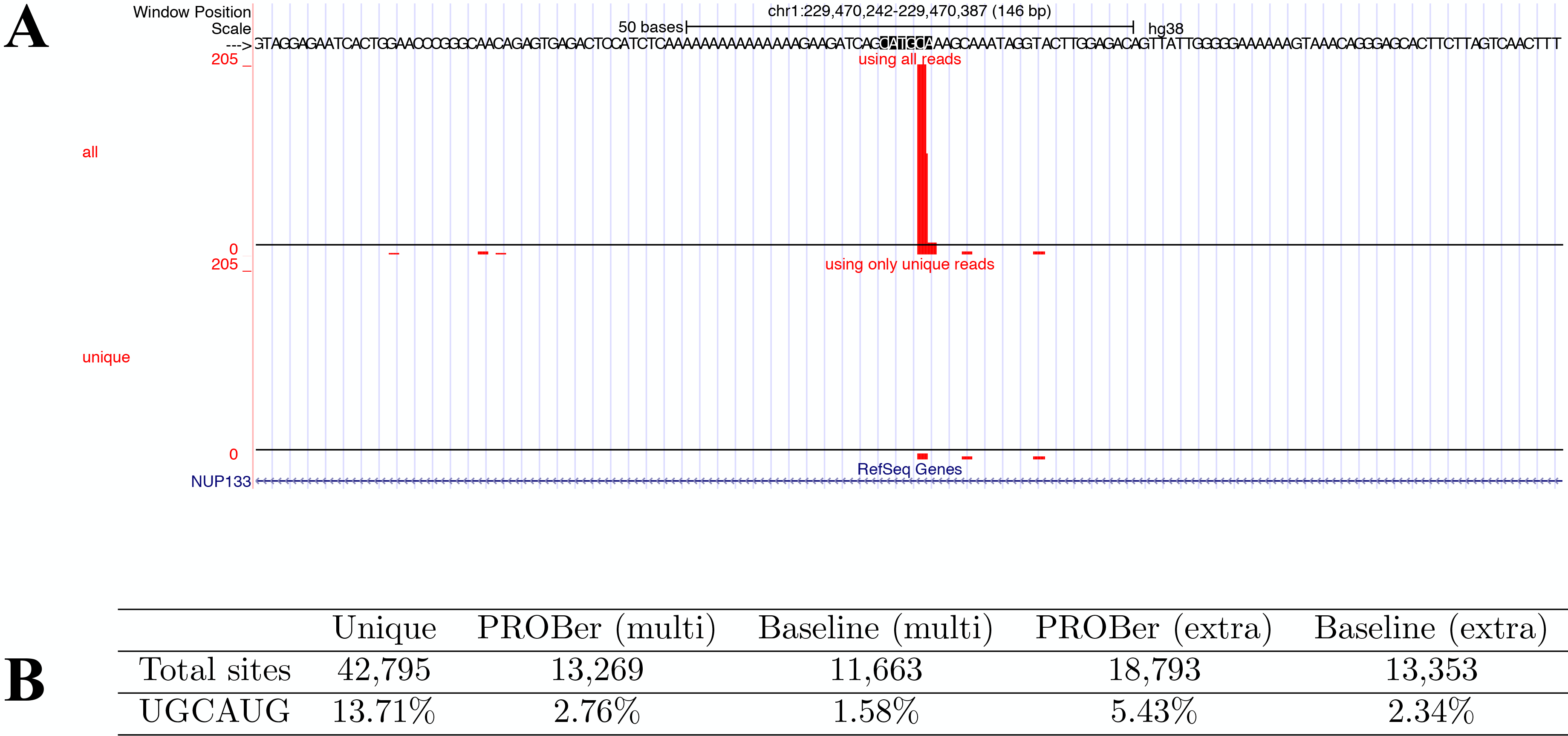
PROBer detected more RBFOX2 binding sites with canonical motifs by utilizing multi-mapping iCLIP reads. (A) The canonical motif containing *NUP133* binding site can only be detected by using both unique and multimapping reads. This plot is generated with UCSC Genome Browser (Kent et al., 2002). Within the plot, the first track shows the number of iCLIP reads (both unique and multi-mapping) dropped-off at each genomic position and the second track only shows the number of unique reads dropped-off at each genomic position. The dropped-off position is one base before the 5' end of an iCLIP read. In addition, the canonical motif is highlighted in the genomic sequences. (B) Numbers of putative RBFOX2 binding sites and the percentage of binding sites that contain the canonical motif UGCAUG. Only sites with at least 10 iCLIP counts are considered as putative binding sites. The canonical motif was searched around each putative binding site using a ±10 nt window. In the first row, Unique refers to binding sites that consist of only uniquely mapped iCLIP reads. PROBer (multi) and Baseline (multi) refers to binding sites that consist of only multi-mapping iCLIP reads and are identified by PROBer and the baseline algorithm respectively. Similarly, PROBer (extra) and Baseline (extra) refers to binding sites that could be identified by using all reads, but not using only uniquely mapped reads. The baseline algorithm distributes each multi-mapping read evenly to its alignments. PROBer identified one times more multi-mapping reads only binding sites with canonical motifs and two times more extra binding sites with canonical motifs than the baseline algorithm.

## DISCUSSION

We present PROBer, a statistically rigorous approach to quantify chemical reactivity profiles from transcriptome-wide sequencing data. We have evaluated PROBer’s performance with three diverse chemical modification protocols, as well as a variety of library preparation protocols. In each of these cases, PROBer outperformed ad hoc methods in analysis of the data. As it is becoming clear that a systems-wide view of such post-transcriptional regulation processes is highly informative, we believe that multiple of these chemical modification / toeprinting protocols will be performed within the same study. As such a unified pipeline such as PROBer is even more valuable.

PROBer is freely available with open-source at http://pachterlab.github.io/PROBer. All experiments can be replicated using the snakemake scripts at https://github.com/pachterlab/PROBer_paper_analysis.

## EXPERIMENTAL PROCEDURES

### Transcript and genome references

*Arabidopsis thaliana*. We downloaded the latest genome and gene annotation (TAIR10) from The Arabidopsis Information Resource. Following Ding et al. (Ding et al., 2014), we extracted every mRNA, rRNA, tRNA, ncRNA, snRNA, miRNA, and snoRNA annotated in the GFF3 file. We also discovered and thus removed 568 duplicate sequences. In addition, we found two copies of 18S rRNA with minor differences and no 25S rRNA (but a subsequence of it, AT2G01021.1) in the extracted sequences. Thus, we added 25S rRNA sequence from the RNA structure database (Cannone et al., 2002) and removed one copy of 18S rRNA, AT3G41768.1, and the 25S subsequence, AT2G01021.1. The final reference consists of 36,264 transcripts in total.

*Saccharomyces cerevisiae*. We downloaded the genome (R64-1-1) and gene annotation (build R64-1-1.84) from Ensembl. After removing duplicate sequences, the final reference consists of 6,841 transcripts.

*Mus musculus*. We downloaded the genome (GRCm38) and gene annotation (build GRCm38.74) from Ensembl. The annotation contains no 18S or 25S rRNAs, and 353 variants of 5S rRNA. We added 18S sequence from the RNA structure database and removed all but one variant of 5S rRNA. We could not add 25S sequence because it is not included in the RNA structure database (Cannone et al., 2002). After removing duplicate sequences, the final reference consists of 93,362 transcripts.

*Homo sapiens*. We downloaded the human genome (GRCh38) from Ensembl.

### Sequencing data

Structure probing data from (Ding et al., 2014) were downloaded from Sequence Read Archive (SRP027216). This data set contains two biological replicates, which were pooled together. We preprocessed the data according to (Ding et al., 2014) which includes removing ssDNA linker and trimming adapter sequence using cutadapt (Martin, 2011) (v1.10). The pre-processed data contain 117,242,295 and 81,596,350 single-end reads in modification-treated and mock-treated experiments respectively. Structure probing data from (Talkish et al., 2014) were downloaded from Sequence Read Archive (SRP029192). Only wild-type data were used and the two biological replicates were pooled together. Data were pre-processed following (Talkish et al., 2014). The pre-processed data contain 7,729,251 and 9,199,721 single-end reads in modification-treated and mock-treated experiments respectively. Structure probing data from (Spitale et al., 2015) were downloaded through Gene Expression Omnibus (GSE64169). The data volume in (Spitale et al., 2015) precluded analysis of the entire dataset so we used only the first 100 million reads from biological replicate 2 of v6.5 mouse ES cells. We pre-processed these data by trimming 3’ adapters, removing PCR duplicates, and then removing unique molecular identifiers (UMI). Only reads with the same sequences and UMIs are considered as duplicates. The data we used consist of three conditions: mock-treated, *in vitro* modification-treated, and *in vivo* modification-treated. After pre-processing, the three conditions contain 96,120,565, 23,455,089, and 78,180,398 single-end reads respectively.

Pseudo-seq data from (Carlile et al., 2014) were downloaded from Gene Expression Omnibus (GSE58200). Following advice of the authors, samples GSM1403085 and GSM1403086 were picked as mock-treated experiments and samples GSM1403087 and GSM1403088 were picked as modification-treated experiments. Data were pre-processed as documented in (Carlile et al., 2014). The resulting pre-processed data contain 31,103,632 and 39,167,224 single-end reads in modification-treated and mock-treated experiments respectively.

RBFOX2 iCLIP data from Nostrand et al. (Van Nostrand et al., 2016) were downloaded at Sequence Read Archive. We only used one run of iCLIP data (SRR3147675). Data were pre-processed by trimming 3’ adapters, removing PCR duplicates, and then removing UMIs. The pre-processed data contain 18,724,388 single-end reads in total.

### PROBer’s generative probabilistic model

We model sequencing-based ‘toeprinting’ experiments using a generative probabilistic model (Figure S3A). The key parameters that we model include the relative abundances for the set of transcripts in the sample, as well as modification probabilities, and RT noise probabilities for each site on a transcript. In order to reduce the number of parameters we have to estimate, we assume the abundances in the modification-treated experiment are the same as abundances in the mock-treated experiment.

To generate a read from the modification-treated experiment, we first pick a transcript at a rate proportional to the product of transcript abundance and length. We denote this rate by *α_i_*. Then we choose the priming site uniformly across all valid priming sites in the transcript. We denote the total number of available priming sites by 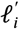. Once we have the priming site, reverse transcription starts. At each site *j*, there is a probability that RT stops due to either chemical modification (denoted by *β*_*ij*_) or background noises such as RT natural drop-off, primer collision or reaching the end of a fragment (denoted by *γ*_*ij*_). Once the RT stops, a cDNA fragment is generated. Thus, the probability of generating a cDNA fragment of length *l*, priming at *j*, and from transcript *i* is

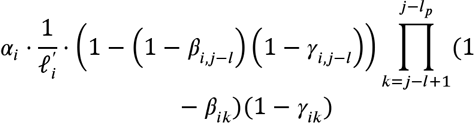

The term *l_p_* in the above equation is the random primer length. In the Ding et al. protocol, this term is equal to 6; however if RNA fragmentation-based protocols are used, this number would be 0.

The next step is to decide if the obtained fragment passes the size selection. If not, this fragment will not be sequenced and therefore considered hidden. Otherwise, a sequence read will be produced according to our sequencing error model. Our model can generate either single-end or paired-end reads and allows both substitution and indel errors to occur during the sequencing step.

To generate a read from the mock-treated experiment is similar, excepting that the chemical modification probabilities are not involved. Thus the probability of generating a similar cDNA fragment becomes

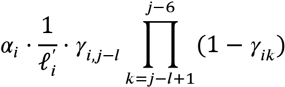

Our generative model is applicable to fragment-based probing protocols (Carlile et al., 2014; Rouskin et al., 2014; Spitale et al., 2015; Talkish et al., 2014) as well. We just need to replace the assumption of uniform priming with the assumption of uniform fragmentation. For more details about our generative model, please refer to Supplemental Experimental Procedures, section 2.

### Estimating PROBer parameters

Our goal is to estimate toeprinting parameters and relative abundances in the sample. Toeprinting parameters include both modification probabilities (*β*s) and RT noise probabilities (*γ*s) per transcript site. We obtain maximum likelihood (ML) estimates for transcript abundances. But for toeprinting parameters, we seek maximum a posteriori (MAP) estimates instead, because for most transcripts, we do not have enough coverage per site to obtain reliable ML estimates. To obtain MAP estimates, we introduce a Beta distribution for each structural parameter (either *β* or *γ*) as its prior. We tie the Beta distribution parameters together for all *β*s and all *γ*s respectively, and set them to 0.0001 by default.

We have two types of hidden data. First, due to alignment ambiguity we cannot be sure about which transcript a read originates from; we can only infer a set of highly possible origins for the read using its alignments. Second, if a cDNA fragment does not pass the size selection, we cannot observe a read from it. For reasons explained in Supplemental Experimental Procedures, section 2.1, we only consider the first type of hidden data.

We use the Expectation-Maximization (EM) algorithm (Dempster et al., 1977) to learn above model parameters. The workflow of our EM algorithm is shown in Figure 1C. In the E step, we interpolate the hidden data– the locations of multi-mapping read – given the estimated abundances and toeprinting parameters. In the M step, we calculate the ML and MAP estimates based on both the observed data and interpolated hidden data. The E and M steps are repeated until convergence. Supplementary Section 3 provides a detailed discussion about our EM algorithm.

### PROBer’s iCLIP model

Because iCLIP protocol does not have a sequenced control and iCLIP signals are sparse in the genome, PROBer only allocates multi-mapping reads for iCLIP data. For this reason, PROBer uses a simpler generative model (Figure S1B). To generate an iCLIP read, PROBer first picks a crosslink site and then generates the read sequence according to a sequencing error model. PROBer uses an Expectation-Maximization-Smoothing (EMS) algorithm (Silverman et al., 1990), which is similar to Chung et al.’s work on ChIP-Seq data (Chung et al., 2011), to infer model parameters and allocate multi-mapping reads. Please refer to Supplemental Experimental Procedures, Section 5 for more details.

### The PROBer software

PROBer contains five commands: prepare, estimate, simulate, iCLIP and version.

The first step in running PROBer is to build reference indices using the command *PROBer prepare*. The command accepts either a genome or a set of transcript sequences. If the input is a genome, users need to specify either a GTF file using the option --gtf or a GFF3 file using the option --gff3. Then PROBer will automatically extract transcript sequences from the specified annotation file. In addition, *PROBer prepare* can help users to build Bowtie (Ben Langmead et al., 2009) and Bowtie 2 (Ben Langmead and Salzberg, 2012) indices by enabling --bowtie and --bowtie2 options. For iCLIP data, --genome option should be set to notify PROBer that genome indices, instead of transcript indices, are required. *PROBer prepare* only needs to be run once per reference.

Next, *PROBer estimate* is run on ‘toeprinting’ data (except iCLIP data). PROBer accepts either raw reads in FASTA/FASTQ format or alignments in SAM/BAM/CRAM format as its inputs. It can handle single-end reads, paired-end reads and indel alignments. If inputs are raw reads, PROBer will call Bowtie to align them by default. Users can ask PROBer to use Bowtie 2 instead by enabling --bowtie2 option. PROBer outputs ML estimates of transcript abundances and MAP estimates of modification and RT noise probabilities. If --output-bam is enabled, PROBer in addition outputs BAM files consisting of posterior-probability-annotated read alignments. PROBer can run with only modification-treated data if mock-treated control is not available. In that case, the estimated modification probabilities might not be as accurate.

*PROBer estimate* options include --primer-length, --size-selection-min, --size-selection-max, and --read-length. --primer-length determines the random primer length. This option should be set to 6 if random hexamer priming was used and to 0 if the protocol was fragmentation-based. --size-selection-min and --size-selection-max describe the minimum and maximum cDNA fragment lengths in your library after size selection. --read-length is only used for single-end reads and specifies the untrimmed read length. It helps PROBer to determine which single-end reads are adaptor trimmed and thus can be regarded as full fragments.

For iCLIP data, we run *PROBer iCLIP*. Similar to other ‘toeprinting’ assays, PROBer accepts iCLIP data either as raw reads in FASTA/FASTQ formator as alignments in SAM/BAM/CRAM format. If inputs are raw reads, either Bowtie or Bowtie 2 can be called to align these reads. However, because of the differences between iCLIP and other ‘toeprinting’ protocols described in the main text, PROBer only allocate multimapping reads for iCLIP data. The PROBer outputs consist of every iCLIP crosslink site and its unique read count & expected multi-read count.

If users want to simulate ‘toeprinting’ reads based on parameters learned from real data, *PROBer simulate* should be used. The simulation parameters can be learned using *PROBer estimate*. Note that PROBer currently cannot simulate iCLIP data.

*PROBer version* prints out the version information.

### Methods used in structure-probing experiments

We compared PROBer with three other methods: StructureFold (Tang et al., 2015), Mod-seq (Talkish et al., 2014), and icSHAPE (Spitale et al., 2015). These three methods were proposed and used in Ding et al. (Ding et al., 2014), Talkish et al. (Talkish et al., 2014), and Spitale et al. (Spitale et al., 2015) respectively. We re-implemented each method according to its original paper. Note that icSHAPE requires a parameter α for the mock-treated experiment. We set α to 0.25, which is the value used in (Spitale et al., 2015) for all structure-probing data sets.

We used Bowtie (Ben Langmead et al., 2009) (v1.1.2) to align reads for all these methods (including PROBer). Because structure-probing protocols are strand-specific, we only aligned reads to the forward strand. For the (Ding et al., 2014) Arabidopsis data, we required at most 3 mismatches for each qualified alignment. For all other data sets, we used Bowtie’s default setting. To allocate multi-mapping reads, PROBer used all qualified alignments of a read. In addition, reads with more than 200 qualified alignments were filtered out. StructureFold and icSHAPE used all qualified alignments in the best stratum (least number of mismatches in either entire read or the “seed” region). Mod-seq used only the best single qualified alignment. These parameter settings were chosen according to the papers describing each method.

PROBer’s protocol-specific options, such as-- primer-length, --size-selection-min, --size-selection-max, and- --read-length, were set differently according to the characteristics of each protocol. Spitale et al. used biotin to selectively enrich structural signals in modification-treated experiments^8^. This step significantly reduces the background noise contained in the modification-treated channel and also makes it hard to interpret the relationship between mock-treated and modification-treated channels. Thus, for Spitale et al. data, we only used modification-treated data as PROBer’s input. For further details, please refer to our Snakemake (Köster and Rahmann, 2012) workflow.

### Simulation of structure-probing experiments and digital spike-in experiments

To assess the variability of the simulation, we simulated two sets of 30 million 37 nt single-end reads in both the modification-treated and mock-treated experiments, using the generative model described before. The model parameters used in the simulation were learned from the Ding et al. structure-probing data by running PROBer. To access if structure information can affect RNA-Seq quantification process (Figure S4), we in addition simulated 30 million 37 nt single-end reads using the RSEM simulator (Li and Dewey, 2011) (which ignores structure information) with the same simulation parameters.

For digital spike-in experiments, our transcriptome was augmented with sequences of three model RNAs from Lucks et al. (Lucks et al., 2011): 1) RNase P from *Bacillus subtilis*, 198 nt; 2) pT181 long from *Staphylococcus aureus*, 192 nt; 3) pT181 short from *Staphylococcus aureus*, 172 nt. 2) and 3) are two variants of the pT181 transcriptional attenuator. The RNA structure and RT noise parameters for these transcripts were calculated from SHAPE-seq data according to Aviran et al. (Aviran et al., 2011b). In order to explore the effect of expression level on estimation accuracy, we generated 4 sets of simulated data by varying the ground truth expression levels of the three RNAs between 100, 1000, 10,000, and 100,000 *Transcripts Per Million* (TPM). Each set of simulated data consists of 30 million 37 nt single-end reads for both the modification-treated and mock-treated experiments. Model parameters for the rest of the transcriptome were set to the same values as described above.

### Comparison with other methods on simulated data

Our main simulation results are box plots comparing PROBer with alternative methods. In these box plots, we only focused on 1,802 transcripts that we may obtain reasonable RNA structure estimates. These transcripts were selected according to the following criteria: 1) its ground truth expression level ≥ 50 TPM; 2) its length ≥ 100 nt, and 3) its mappability score > 0. The mappability score is defined as the ratio between the number of 21 mers that can be mapped back uniquely and the total number of 21 mers in the same transcript. We further partitioned the 1,802 transcripts into 4 expression ranges in TPM: 887 transcripts in [50, 100], 849 transcripts in (10^2^, 10^3^], 60 transcripts in (10^3^, 10^4^], and 6 transcripts in (10^4^,10^6^].

For each transcript and method, we calculated Pearson’s correlation coefficient between the ground truth modification probabilities and the estimates. In the calculation, we only used sites containing ‘A’s or ‘C’s because DMS only modifies ‘A’s and ‘C’s. In addition, we excluded the last 36 nt (read length is 37 nt) of each transcript from the analysis because there are little reads aligned to the 3’ end.

In addition to the results shown in Figure2A, we also investigated the effects of interpolating hidden fragments that failed to pass size selection. We named PROBer with this interpolation enabled as the full model (see Supplemental Experimental Procedures, section 3). As shown in Figure S2A, the full model significantly increased the variance for structural estimates in medium expression ranges, which contain over 96% of investigated transcripts. This result validates our decision of taking off the size selection correction step from PROBer. To demonstrate to the improvement in performing structure estimation and transcript quantification at the same time, we also compared PROBer with the RSEM + PROBer* pipeline. RSEM (Li et al., 2010; Li and Dewey, 2011) is a popular RNA-Seq transcript quantification software that is not aware of RNA structure information. PROBer* is a modified version of PROBer that only works on a single transcript and thus is not aware of multi-mapping reads. Figure S2B confirms our hypothesis --- PROBer performs better at all expression ranges than the RSEM+PROBer* pipeline.

To assess the variability introduced by simulation, we simulated an extra data set. Boxplots for this simulation (Figure S3), demonstrate that our results are stable with respect to the simulation used.

### Comparison with other methods using ROC curves

We compared PROBer’s MAP estimates of chemical modification probabilities with alternative methods’ scores using previously reported ribosomal RNA secondary structures (Cannone et al., 2002). Secondary structures for Arabidopsis 18S and 25S rRNAs, yeast 18S and 25S rRNAs, and mouse 18S and 12S mitochondrial rRNAs were obtained as BPSeq files. Sites on these rRNAs that participate in a base-pairing interaction were assigned an idealized modification rate of 0, and unpaired sites were assigned an idealized modification rate of 1. ROC curves comparing PROBer estimated MAP chemical modification rate and alternative method scores to this binary ground truth vector were produced and the areas under the ROC curves were calculated using PRROC (Keilwagen et al., 2014).

### Experiments on Carlile et al. Pseudo-seq data

For yeast, we have 49 known Ψ sites. However, the rRNAs and snoRNA containing these Ψs have 1905 thymines (T). Thus, this data set is highly skewed. It is known that when data sets are highly skewed, ROC curves tend to be overly optimistic (Davis and Goadrich, 2006). In fact, we can observe this phenomenon by plotting the ROC curves of this data set (Figure S7). Thus, in the main text, we chose Precision-Recall (PR) curve to evaluate different methods. PR curves were produced using PRROC (Keilwagen et al., 2014). In addition, we have observed a strange read count pattern at the 5’ end of 25S rRNA. Normally, the 5’ end base of a transcript should have a very high read count because of RT run-off. However, for 25S, the high read count appears at the 3rd base. We hypothesize that this may be due to a small amount of degradation in the input RNA.

### Reproducing our experiments

We implemented a Snakemake (Köster and Rahmann, 2012) workflow which can be used to replicate all our analyses: https://github.com/pachterlab/PROBer_paper_analysis

## AUTHOR CONTRIBUTIONS

Development of statistical framework for toeprinting assays and formulation of the PROBer model: B.L., A.T., S.A., and L.P.; Software and analysis: B.L.; Preparation fo manuscript: B.L., A.T., S.A., and L.P.; Funding Acquisition, L.P.; Supervision, S.A. and L.P.

## ACKNOWLEDGMENTS

We thank Yiliang Ding, Joel McManus, and Thomas Carlile for discussions and clarifications on the structure-seq, Mod-seq, and Pseudo-seq methods respectively. We thank Yeon Lee and Julian König for discussions on the iCLIP method. This work is supported by National Institutes of Health (NIH) grants R01 HG006129 to L.P. and R00 HG006860 to S.A., and by the Center for RNA Systems Biology at UC Berkeley (NIH P50GM102706 grant). A.T. was partially supported by NIH Molecular Biophysics Training grant (NIH GM08295).

